# Mutualist-pathogen co-colonisation modulates phosphoinositide signatures at host intracellular interfaces

**DOI:** 10.1101/2025.06.23.661106

**Authors:** Alex Guyon, Theresa Staps, Lyne Badot, Sebastian Schornack

## Abstract

The host membrane that surrounds intracellular microbes forms a critical interface influencing whether interactions result in mutualism or pathogenesis. While phosphoinositide identities differ between pathogen and mutualist interface membranes, it is unclear if these are modulated during co-colonisation. To address this, we generated *Nicotiana benthamiana* plants expressing biosensors for PI4P and PI(4,5)P_2_ and imaged root colonisation by the pathogenic oomycete *Phytophthora palmivora* and the mutualistic fungus *Funneliformis mosseae*. Binary host-microbe interactions revealed distinct patterns: PI(4,5)P_2_ was tip-enriched at mutualist structures but evenly distributed around pathogen structures, while PI4P was absent from pathogen-associated membranes but present at mutualist interfaces. Strikingly, co-colonisation altered host membrane identity, triggering PI4P recruitment at pathogen haustoria, and enhanced resistance to *P. palmivora*. These findings reveal that phosphoinositide signatures distinguish pathogenic and mutualistic interfaces and are dynamically remodelled during co-colonisation, likely influencing interaction outcomes.

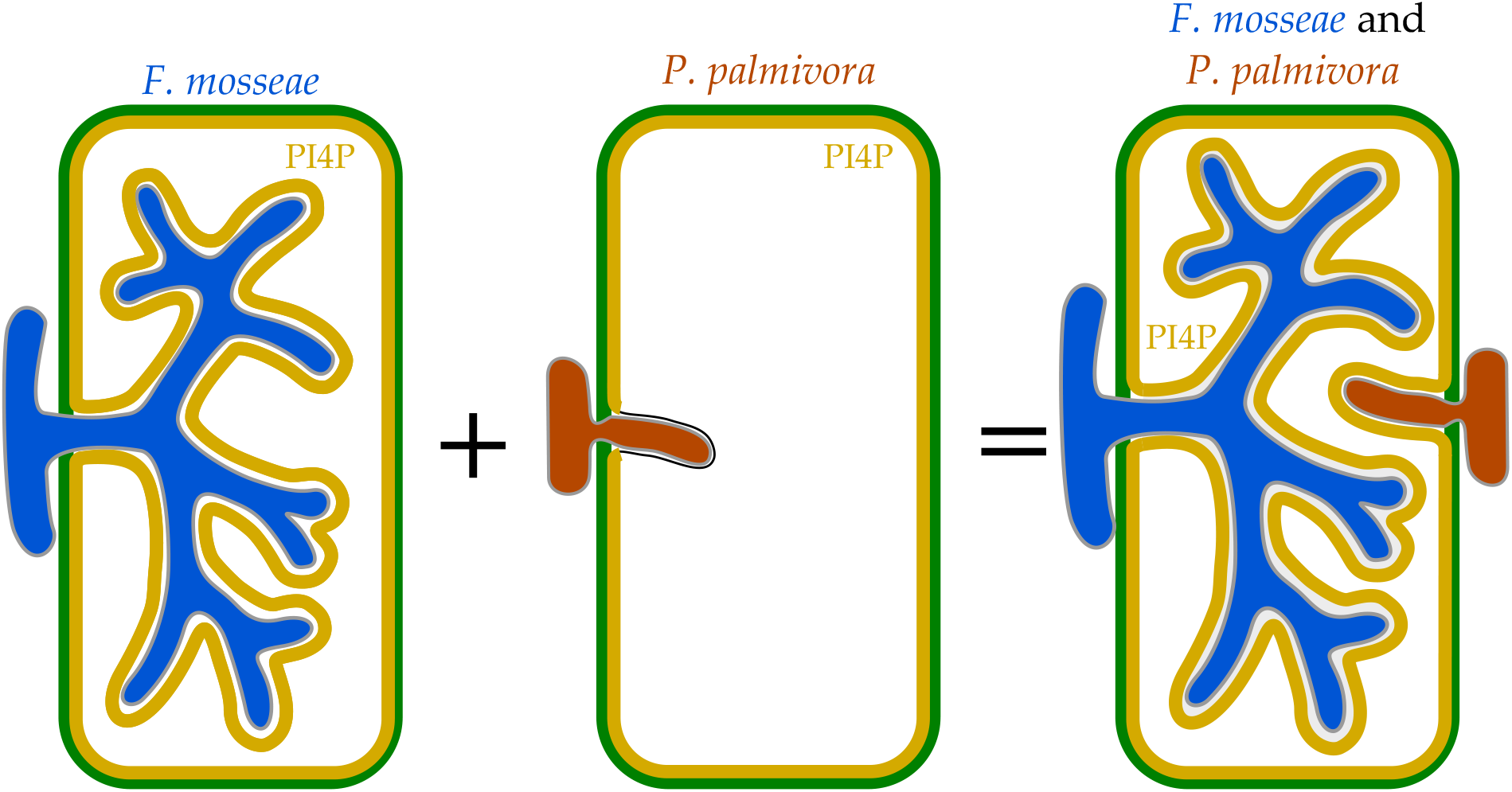

## INTRODUCTION

Plants interact with a range of microbes, both above and below ground. These interactions span a spectrum from mutualism (where both partners benefit) to parasitism (where the microbe benefits at the expense of the host) (Hirsch, 2004). One of the most ancient and widespread mutualisms is the arbuscular mycorrhizal (AM) symbiosis, occurring between most land plants and Glomeromycotina fungi such as *Funneliformis mosseae* (Smith and Read, 2008). In contrast, microbes can act as pathogens that reduce fitness or cause death of their host, including filamentous eukaryotes like the oomycete *Phytophthora palmivora* (Hardham, 2001). Each of these filamentous microbes penetrate the plant cell wall and form intracellular structures that remain separated from the host cytoplasm by specialised, host derived membranes. These include the periarbuscular membrane (PAM) surrounding arbuscules in AM symbiosis (Pumplin and Harrison, 2009), and the extrahaustorial membrane (EHM) surrounding haustoria formed by pathogens (Lu et al., 2012). These interface membranes are not passive barriers; they are essential sites for nutrient exchange and trans-kingdom communication. The PAM is the site of lipid transfer from host to AM fungus via *STR/STR2* transporters, and phosphate uptake via the PT4 transporter (Harrison et al., 2002; Jiang et al., 2017; Luginbuehl et al., 2017; Pumplin and Harrison, 2009). Pathogens, in turn, secrete host-manipulating microbial proteins, termed effectors, that can be delivered into host cells via haustoria, crossing the EHM to modulate intracellular processes and immune responses (Bozkurt et al., 2012; Wang et al., 2017).

Although the PAM and EHM are both continuous with the host plasma membrane, they have distinct identities, both in protein and lipid composition, that impact interaction outcomes (Lu et al., 2012). Phosphoinositides (also called phosphatidylinositol phosphates, PIPs) are prominent lipids contributing to membrane identity in plant cells (Noack and Jaillais, 2017). These are a conserved family of phosphorylated derivatives of the membrane phospholipid phosphatidylinositol, such as PI4P and PI(4,5)P_2_, that are distributed in specific patterns across endomembrane compartments and are recognised by protein domains that specifically bind different phosphoinositide species. In AM symbiosis of *Medicago truncatula*, PI4P is found at the periarbuscular membrane, while PI(4,5)P_2_ is enriched at specific subdomains of the PAM, showing a more dynamic distribution (Ivanov and Harrison, 2019). In contrast, in interactions with filamentous pathogens forming haustoria, PI(4,5)P_2_ is present at the EHM, whereas PI4P is consistently excluded (Qin et al., 2020; Shimada et al., 2019). These patterns suggest that distinct phosphoinositide signatures mark mutualistic versus pathogenic interfaces possibly influencing their functions.

While plant–microbe interactions are often studied in simplified binary systems, plants in nature are colonised by complex microbial communities, and even individual cells can host multiple microbes. For example, Belhaj et al., 2017 showed that *Arabidopsis* cells can simultaneously accommodate the oomycetes *Phytophthora infestans* and *Albugo candida*. However, whether a single plant cell can simultaneously accommodate both mutualist and pathogenic structures, and how this impacts the identities of their respective interface membranes, remains unclear.

To investigate how interface membrane identity is shaped during co-colonisation, we generated stable *Nicotiana benthamiana* lines expressing biosensors for the phosphoinositides PI4P and PI(4,5)P2. Using these lines, we characterised the interface membranes formed during both binary and co-colonisation plant-microbe interactions with pathogenic (*P. palmivora*) and mutualistic (AM fungus *F. mosseae*) microbes. We discovered that PI(4,5)P2 was uniquely tip-enriched around mutualist structures but evenly distributed around pathogen structures. By contrast, PI4P was largely excluded from extrahaustorial membranes but present at periarbuscular membranes of the mutualist fungus. Significantly, mutualistic pre-colonisation enhanced resistance to *P. palmivora* and gave rise to root cells simultaneously hosting mutualistic arbuscules and pathogenic haustoria. Strikingly, haustoria in mycorrhizal roots possessed a distinct membrane identity by recruiting PI4P, demonstrating the plasticity of interface membrane identity.

## RESULTS

### PI(4,5)P_2_ marks host membranes around microbial structures with enrichment at AM fungal tips

To determine if PI(4,5)P_2_ distinguishes membranes interfacing with pathogenic versus mutualistic microbes, we generated a stable transgenic *Nicotiana benthamiana* line expressing the PIPline biosensor P24Y (*Citrine-2xPH*^*PLC*^), which specifically binds phosphatidylinositol 4,5-bisphosphate (PI(4,5)P_2_). This sensor had previously been characterised in *Arabidopsis thaliana*, a non-host for arbuscular mycorrhizal (AM) fungi (Simon et al., 2014). P24Y is a high avidity biosensor, featuring tandem PH^PLC^ lipid-binding domains that preferentially mark membranes enriched in PI(4,5)P_2_ due to the cumulative binding strength of multiple interactions. In the absence of microbial colonisation, P24Y localised to both the plasma membrane and cytosol of root epidermal cells (Fig. 1A) with increased membrane localisation upon brief heat stress similar to previous observations in *Nicotiana* BY-2 cells (Mishkind et al., 2009). In leaf epidermal cells, P24Y predominantly marked the plasma membrane, both in stable lines and transient expression assays (Fig. S1A-B).

**Figure 1).**
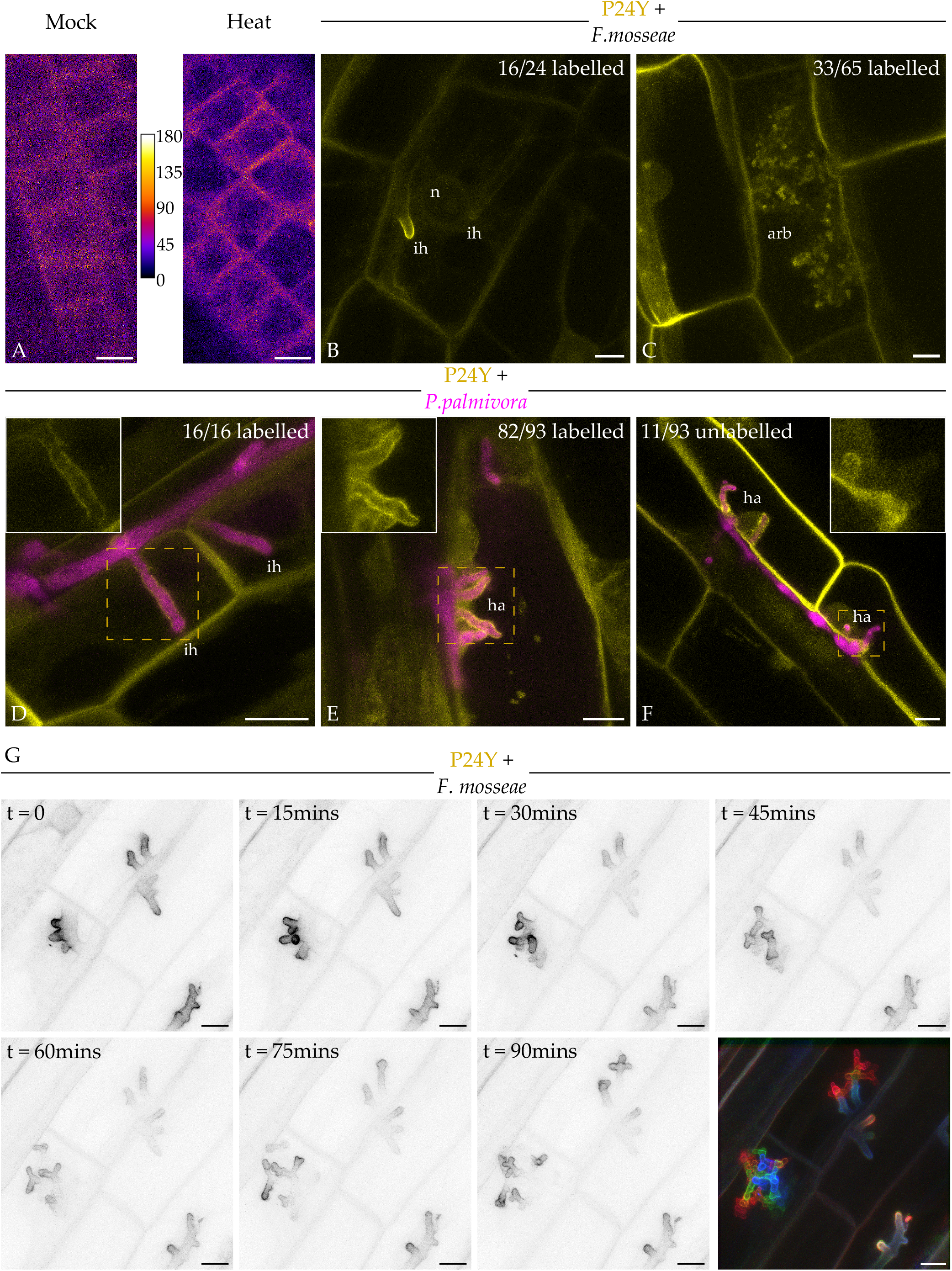
PI(4,5)P_2_ biosensor P24Y labels plant membranes around intracellular *P. palmivora* hyphae and haustoria and accumulates at the tips of growing fungal structures. Confocal imaging of *N. benthamiana* stable transgenic roots expressing the PI(4,5)P_2_ biosensor P24Y (pAtUBQ10::*Citrine-2xPH*^*PLC*^). **A:** Single channel Citrine images (false coloured with indicated look up table) showing P24Y signal under mock conditions or after a 30-min 40°C heat treatment. **B-C:** Roots colonised with *F. mosseae*; images show Citrine channel alone (yellow). **D-F:** Roots infected with *P. palmivora* strain ARI-tdTomato. Images are composites of Citrine (yellow) and tdTomato (magenta) channels. **G:** Time-lapse imaging of a root section colonised by *F. mosseae*, captured over 90 minutes. Panels show inverted grayscale Citrine signal; the final panel is a false-coloured composite of the individual panels indicating progression over time. Images in A are single z-slices, all other images are maximum intensity z-projections. ih, intracellular hyphae; ha, haustoria; n, nucleus; arb, arbuscule. Scale bars = 10µm.

We then challenged *N. benthamiana* lines expressing the PI(4,5)P_2_ biosensor with either the mutualistic AM fungus *Funneliformis mosseae* or the pathogenic oomycete *Phytophthora palmivora* and examined PI(4,5)P_2_ distribution around the structures formed by these microbes within host cells. We found that the PI(4,5)P_2_ biosensor P24Y labelled root-intracellular hyphae of both *F. mosseae* and *P. palmivora* (Fig. 1B, D). Strikingly, P24Y localised specifically to the hyphal tips of *F. mosseae* in two-thirds of observed cases (Fig. 1B), whereas it consistently showed a uniform distribution around *P. palmivora* hyphae (Fig. 1D). Similarly, P24Y labelled arbuscules in roots (Fig. 1C) and was preferentially enriched at their tips in about half of the observed cases, consistent with tip-localised P24Y signal around *F. mosseae* hyphae. In contrast, uniform P24Y signal was observed around the majority of *P. palmivora* haustoria in roots, with an enrichment of signal relative to background cytosolic signal (Fig. 1E). Similarly, a majority of *P. palmivora* haustoria in leaves were labelled by stably expressed P24Y (Fig. S1C).

Given the tip-specific enrichment around mutualist structures, we hypothesised that PI(4,5)P_2_ may accumulate at the growing tips of intracellular fungal structures. To test this, we performed time-lapse imaging of *F. mosseae* colonised roots expressing P24Y using the AMSlide, a non-invasive, high-resolution, live-imaging system (McGaley et al., 2024). We observed membrane-associated P24Y signal that persisted at the tips of intracellular fungal structures as they expanded over time (Fig. 1G, Video S1), while non-growing tips lacked a prominent focal P24Y signal.

In summary, our results reveal that PI(4,5)P_2_ is a component of the extrahaustorial membrane (EHM) around *P. palmivora* haustoria, the extra-invasive hyphal membrane (EIHM) around *P. palmivora* hyphae and the periarbuscular membrane (PAM) around *F. mosseae* arbuscules and hyphae. Additionally, PI(4,5)P_2_ is specifically enriched at the tips of *F. mosseae* arbuscules and intracellular hyphae, suggesting the PAM at these sites possesses a unique identity.

### PI4P is present in the periarbuscular membrane but absent from the extrahaustorial membrane

Similarly, to determine if PI4P distinguishes membranes interfacing with pathogenic versus mutualistic microbes, we generated a stable transgenic *Nicotiana benthamiana* line expressing a different PIPline biosensor P21Y (*Citrine-2xPH*^*FAPP1*^), which specifically binds phosphatidylinositol 4-phosphate (PI4P) (Simon et al., 2014). P21Y is a high avidity biosensor, featuring tandem PH^FAPP1^ lipid-binding domains that preferentially mark membranes enriched in PI4P due to the cumulative binding strength of multiple interactions. In the absence of microbial colonisation, P21Y exhibited strong plasma membrane localisation in roots and leaves (Fig. 2A, S2A, D).

**Figure 2).**
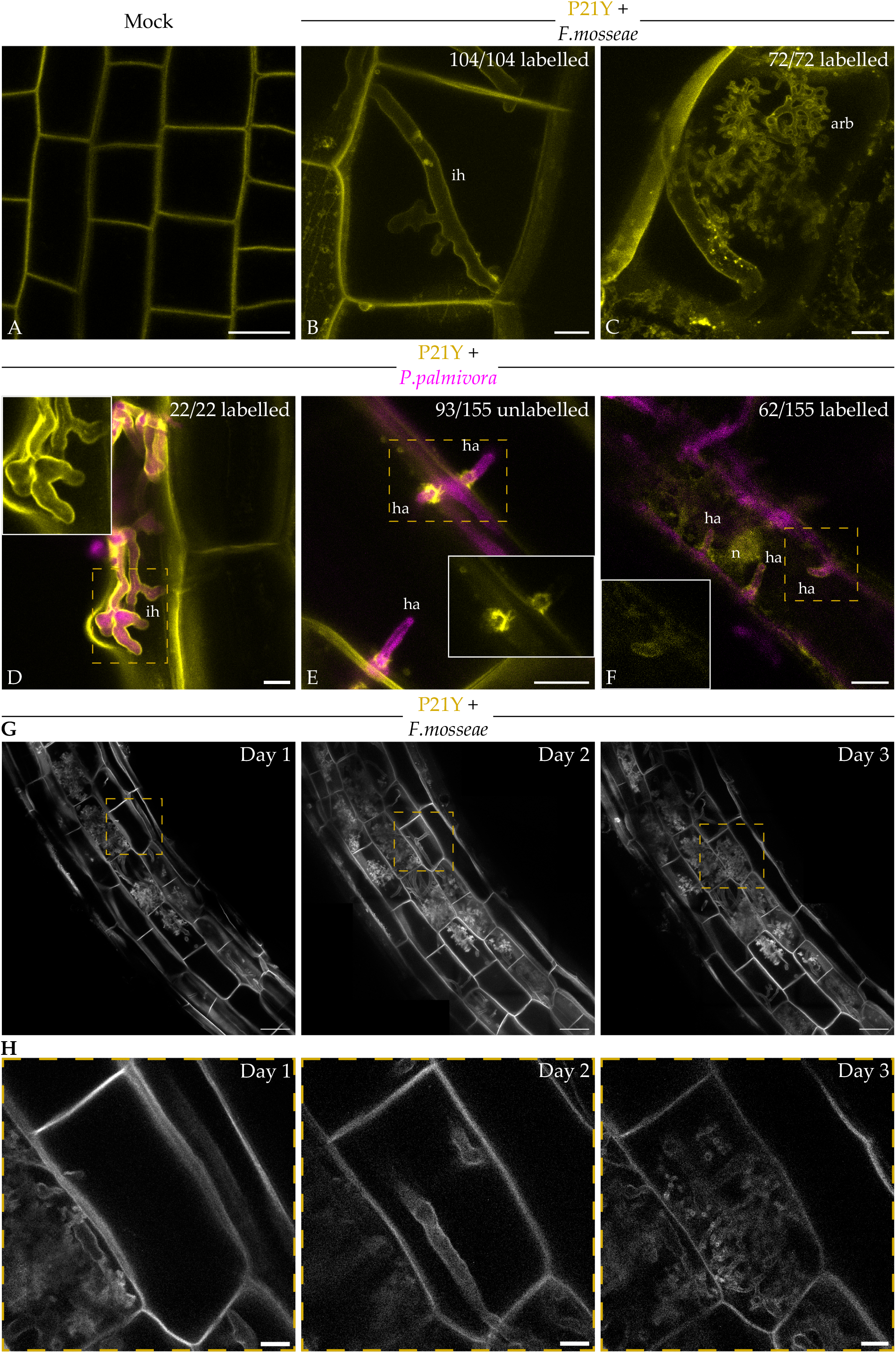
PI4P biosensor P21Y labels arbuscules and hyphae of *F. mosseae* and *P. palmivora* but not haustoria. Confocal imaging of *N. benthamiana* stable transgenic roots expressing the PI4P biosensor P21Y (pAtUBQ10::*Citrine-2xPH*^*FAPP1*^). **A:** P21Y signal in root tip epidermal cells, image shows Citrine channel alone (yellow). **B-C:** Roots colonised with *F. mosseae*, images show Citrine channel alone (yellow). **D-F:** Roots infected with *P. palmivora* strain ARI-tdTomato. Images are composites of Citrine (yellow) and tdTomato (magenta) channels. **G-H:** Time-lapse imaging of a root section colonised by *F. mosseae*, captured over 3 days. Panels show grayscale Citrine signal. **H:** Shows a close-up of the area marked in G. Images A and F are single z-slices, all other images are maximum intensity z-projections. ih, intracellular hyphae; ha, haustoria; n, nucleus; arb, arbuscule. Scales bars in A-F, H = 10µm, in G = 50µm.

We then challenged *N. benthamiana* lines expressing the PI4P biosensor with either the mutualist *F. mosseae* or the pathogen *P. palmivora* and examined PI4P distribution around the structures formed by these microbes within host cells. We found that the PI4P biosensor P21Y always labelled intracellular hyphae of *F. mosseae* or *P. palmivora* (Fig. 2B, D). Additionally, P21Y always clearly labelled arbuscules, with signal tightly and uniformly surrounding the intracellular structures, consistent with localisation at the PAM (Fig. 2C). In contrast to arbuscules, the majority (60%) of *P. palmivora* haustoria in roots did not show P21Y signal (Fig. 2E). Similarly, P21Y was excluded from the majority (69%) of haustoria when transiently expressed in leaves (Fig. S2B). Likewise, in leaves of stably expressing PI4P biosensor plants, a subset of haustoria (26%) lacked detectable PI4P at the EHM with the exception of a labelled neck region (Fig. S2E). However, 74% of haustoria showed P21Y signal fully surrounding them, specifically in cells which also displayed cytosolic biosensor signal around chloroplasts (see inset in Fig. S2F).

We next carried out timelapse imaging of *F. mosseae* colonised roots using the AMSlide (McGaley et al., 2024) to further investigate the localisation of P21Y over time. This revealed uniform localisation of P21Y around all fungal structures observed at all stages (Fig. 2G, H). These results indicate that PI4P is consistently present in the PAM and in membranes around intracellular hyphae of both *F. mosseae* and *P. palmivora* but is excluded from the EHM around a majority of *P. palmivora* haustoria, indicating distinct phosphoinositide identities between host-derived intracellular membranes.

### Haustoria in mycorrhizal roots have a different EHM identity than those in non-mycorrhizal roots

Given the differential distribution of phosphoinositide reporters for PI4P around haustoria and arbuscules, we tested whether these patterns change during co-colonisation of *N. benthamiana* roots with *P. palmivora* and *F. mosseae*. We first colonised roots stably expressing P21Y with *F. mosseae* (+AMF) or grew them with heat-killed *F. mosseae* inoculum as a control (-AMF), then subsequently infected them with *P. palmivora*. As previously observed, most haustoria (82%) formed in the absence of *F. mosseae* colonisation were not labelled by P21Y (Fig. 3A). However, in the presence of *F. mosseae* colonisation, a majority of *P. palmivora* haustoria (83%) were now labelled by P21Y while P21Y localisation remained unchanged around AM fungal structures (Fig. 3B). This shift in EHM identity occurred both in cells that were co-colonised by *F. mosseae* and *P. palmivora* as well as in neighbouring cells without arbuscules but with haustoria.

**Figure 3).**
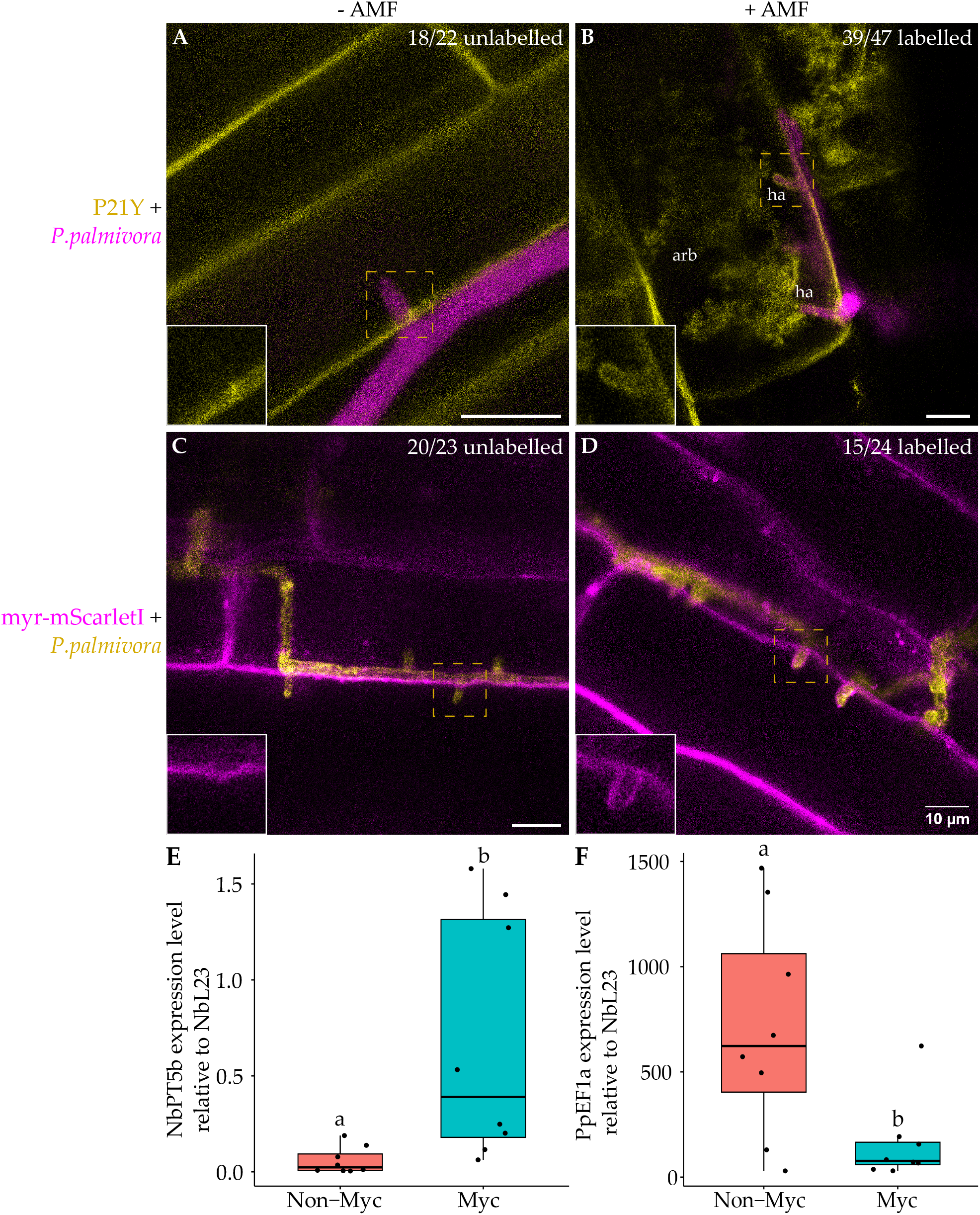
Haustoria in mycorrhizal roots have a different EHM identity than those in non-mycorrhizal roots. **A-B:** *N. benthamiana* stable transgenic roots expressing the PI4P biosensor P21Y (pAtUBQ10::*Citrine-2xPH*^*FAPP1*^) infected with *P. palmivora* strain ARI-tdTomato in the absence (**A**) or presence (**B**) of AM colonisation. Images are composites of the Citrine channel (yellow) and the tdTomato channel (magenta). **C-D:** *N. benthamiana* stable transgenic roots expressing pAtUBQ10::*myr-mScarletI* infected with *P. palmivora* strain LILI-YKDEL in the absence (C) or presence (D) of AM colonisation. Images are composites of the YFP channel (yellow) and the mScarletI channel (magenta). Images A & D are maximum intensity z projections, other images are single z slices. ha, haustoria; arb, arbuscule. Scale bars = 10µm. **E-F:** RT-qPCR data of *NbPT5b* and *PpEF1α* respectively during infection of mycorrhizal or non-mycorrhizal *N. benthamiana* roots by *P. palmivora* ARI-tdTomato. These trends were seen in 2/3 replicates, statistical groupings are from a Kruskal-Wallis test, p < 0.05, n = 8.

To determine whether changes in the EHM extend beyond PI4P, we examined the distribution of a myristoylated mScarletI fluorescent protein (myr-mScarletI) which is targeted to membranes via a myristic acid lipid anchor. Interestingly, myr-mScarletI was excluded from membranes surrounding haustoria in non-mycorrhizal roots (Fig. 3C) but was present around haustoria in mycorrhizal roots (Fig. 3D).

Because the shift in EHM identity appeared to occur at a tissue level, we tested whether it could be systemic. We grew plants expressing myr-mScarletI in the presence or absence of *F. mosseae* colonisation and their leaves were subsequently infected with *P. palmivora*. In both conditions, myr-mScarletI was absent from around the majority of haustoria (Fig. S3A, B) suggesting the effect is restricted to root tissue and not systemically induced.

To test whether haustoria formed in mycorrhizal roots are viable we used the *P. palmivora* strain LILI *Avr3a-mRFP*, which expresses an effector from *P. infestans* (Whisson et al., 2007). This effector is secreted from haustoria and accumulates in the extrahaustorial matrix, resulting in a specific enrichment of mRFP signal around haustoria compared to apoplastic hyphae. We observed AVR3a-mRFP signal around haustoria in both mycorrhizal and non-mycorrhizal roots (Fig. S3C, D). This suggests that haustoria forming in mycorrhizal roots are still able to secrete effectors to the host-pathogen interface. No AVR3a-mRFP signal was detected inside plant cells, even in non-mycorrhizal roots, suggesting that this fusion protein is not taken up in detectable amounts.

To test whether changes in phosphoinositide membrane identity correlate with altered pathogen infection, we examined whether *F. mosseae* colonisation affects the extent of *P. palmivora* colonisation. The AM marker gene *NbPT5b*, which is specifically expressed in arbuscule-containing cells was significantly upregulated in *F. mosseae* colonised roots (Myc) compared to controls (Non-Myc), confirming successful mycorrhizal colonisation (Fig. 3E). The expression of the *P. palmivora* biomass marker *PpEF1a* was significantly lower in mycorrhizal roots, indicating reduced pathogen colonisation (Fig. 3F).

In summary, the phosphoinositide identity of membranes around intracellular microbes is dynamic and can be modulated by simultaneous colonisation with another microbe, and these phospholipid biosensor dynamics have not been reported before, likely due to the binary nature of most plant-microbe studies.

## DISCUSSION

Our study reveals that the identities of plant-microbe interfaces are not static but can be dynamically reshaped by co-colonisation with mutualistic and pathogenic microbes. Using phosphoinositide biosensors in *Nicotiana benthamiana*, we show that the distribution of PI4P and PI(4,5)P_2_ distinguishes mutualistic and pathogenic interfaces in roots, and that these signatures are altered in the context of co-colonisation.

We discovered that PI(4,5)P_2_ specifically labels the plant membrane regions around the growing tips of intracellular AM fungal structures, likely reflecting a role in polarised deployment of new membrane material at these sites and providing a valuable marker facilitating time-lapse imaging of AM colonisation. In contrast, PI4P is a more general signature of intracellular colonisation that equally labels the plant membranes surrounding mutualist structures. Tip-localised PI(4,5)P_2_ echoes its role in other polar-growth systems (root hairs, pollen tubes, rhizobia infection threads) suggesting perifungal membrane expansion relies on a conserved polarised secretion machinery (Lace et al., 2023; Noack and Jaillais, 2017). However, this differs from *Medicago truncatula* where tip-localised PI(4,5)P_2_ was observed only at short linear hyphae and newly forming thick branches but never at fine branches of arbuscules of *Rhizophagus irregularis, Glomus versiforme* and *Gigaspora gigantea* (Ivanov and Harrison, 2019). Given the same phosphoinositide binding domain was used in the PI(4,5)P_2_ biosensor in *M. truncatula*, we hypothesise this reflects host or fungal species-specific differences, perhaps due to different colonisation morphologies. *N. benthamiana* exhibits intermediate Arum-Paris type colonisation where arbuscules often branch off intracellular hyphae, whereas *M. truncatula* has Arum-type colonisation with arbuscules forming from a single cell entry point from a branch of an intercellular hyphae (Dickson, 2004).

We showed that PI4P was frequently excluded from the extrahaustorial membrane surrounding *P. palmivora* haustoria, consistent with reports from *Arabidopsis* leaves showing PI4P was absent from the haustoria of *Erysiphe cichoracearum, Albugo candida*, and *Hyaloperonospora arabidopsidis* (Qin et al., 2020; Shimada et al., 2019). This suggests that PI4P exclusion may represent a conserved feature of haustoria interfaces for a diverse range of biotrophic filamentous pathogens in multiple host species and organs (leaves and roots). However, we found that PI4P was detectable at the EHM in a subset of cells, indicating that exclusion is not absolute and may reflect distinct haustorial populations previously documented in hemibiotrophic *Phytophthora* pathogens (Bozkurt et al., 2011). By contrast, PI4P was present around intracellular *P. palmivora* hyphae formed in root tissue (Hardham, 2001), consistent with the presence of PI4P at the extra-invasive hyphal membrane (EIHM) surrounding the biotrophic invasive hyphae of *Colletotrichum higginsianum* (Qin et al., 2020; Shimada et al., 2019). Intracellular *P. palmivora* hyphae can adopt either linear or branched forms, indicating morphological diversity during colonisation.

We therefore propose that *P. palmivora* forms two types of intracellular structures surrounded by plant derived membranes with distinct phosphoinositide identities: hyphae surrounded by an EIHM and haustoria surrounded by an EHM.

Significantly, we revealed that pre-colonisation by a mutualist rendered roots more resistant to *P. palmivora* and strikingly recruited PI4P and a myristoylated fluorophore to newly formed EHMs, indicating a change in membrane identity. Notably, we observed haustoria and arbuscules within the same host cell, highlighting the host’s capability to simultaneously accommodate both mutualistic and pathogenic intracellular structures. Since haustoria morphology and their effector secretion appeared unchanged in mycorrhizal roots and because PI4P is present around intracellular hyphae, PI4P exclusion does not appear to be an intrinsic property of the haustorial interface. Instead, we propose that PI4P exclusion from the EHM reflects a pathogen driven strategy, likely mediated by effectors that manipulate the phosphoinositide identity of the interface membrane. Possible mechanisms include a direct inhibition of plant PI4-kinases (PI4Ks) responsible for PI4P biosynthesis (Noack et al., 2022); exclusion of PI4Ks from the EHM through interference with vesicular transport; or recruitment of host PI4P phosphatases to degrade PI4P at the interface. Although not reported to date in plant-microbe interactions, some effectors could themselves act enzymatically to degrade PI4P at the interface. Analogous effector-mediated phosphoinositide reprogramming has been described in animal systems, where *Legionella pneumophila* secretes effectors that modify the phosphoinositide composition of the host vacuole to promote intracellular survival and replication (Dong et al., 2016; Pike et al., 2019).

Exclusion of PI4P from the EHM likely impacts plant immune signalling as several key immune components depend on PI4P for proper membrane localisation and function. These include intracellular helper nucleotide-binding leucine rich repeat proteins involved in effector triggered immunity and peripheral receptors for pathogen associated molecular patterns, along with their scaffold proteins like remorins (Gronnier et al., 2017; Lu et al., 2012; Saile et al., 2021). In contrast, intracellular mutualist structures, such as arbuscules, rely on the correct localisation of symbiosis-associated receptors (Roth et al., 2018), perhaps justifying the presence of PI4P at the periarbuscular membrane (PAM) and along intracellular fungal hyphae. Thus, in a situation where AM symbiosis enforces interface trafficking (Harrison, 2012; Pumplin et al., 2012) to arbuscules this process seems to overrule any pathogen efforts to exclude PI4P and indirectly immune signalling components from its haustoria.

Plant-microbe interactions are typically studied in simplified binary systems, but these do not reflect the complex microbial communities which plants encounter in nature. Our results demonstrate that co-colonisation can reshape host membrane identity and potentially shift the outcome of interactions. In particular, the recruitment of PI4P to haustorial interfaces in mycorrhizal roots suggests that PI4P exclusion is not an intrinsic feature of haustoria but a dynamic, pathogen driven strategy that can be disrupted. Future work should focus on identifying the molecular mechanisms underpinning this exclusion. Uncovering how these membrane identities are established and manipulated will help reveal how plants coordinate responses to multiple microbes, offering a more realistic view of plant immunity and symbiosis.

### RESOURCE AVAILABILITY

Further information and requests for resources and reagents should be directed to and will be fulfilled by the lead contact, Sebastian Schornack.

## ACKNOWLEDGMENTS

Funding was provided by the School of Biological Sciences Doctoral Training Programme to A.G (G108721), and by the Royal Society (UF160413) and Gatsby Charitable Foundation (G120503) to S.S. We are grateful to Philip Carella for critical reading of the manuscript, and to Jen McGaley for AMSlide blueprints and valuable advice on time-lapse imaging. We thank Matthew Smoker (TSL) for the transformation of LILI *Avr3a-mRFP*. We also acknowledge Matthew Macleod for technical support in maintaining *P. palmivora* cultures and *N. benthamiana* plants.

## AUTHOR CONTRIBUTIONS

A.G. and S.S. designed research, A.G., T.S and L.B. carried out research and analysed data. A.G. prepared figures. A.G. and S.S. wrote the paper.

## SUPPLEMENTAL INFORMATION

**Document S1. Figures S1-S3 and Tables S1-S3**

**Video S1. Composite time-lapse video of membrane-associated P24Y signal at the tips of intracellular fungal structures as they expanded over time**

## MATERIALS AND METHODS KEY RESOURCES TABLE

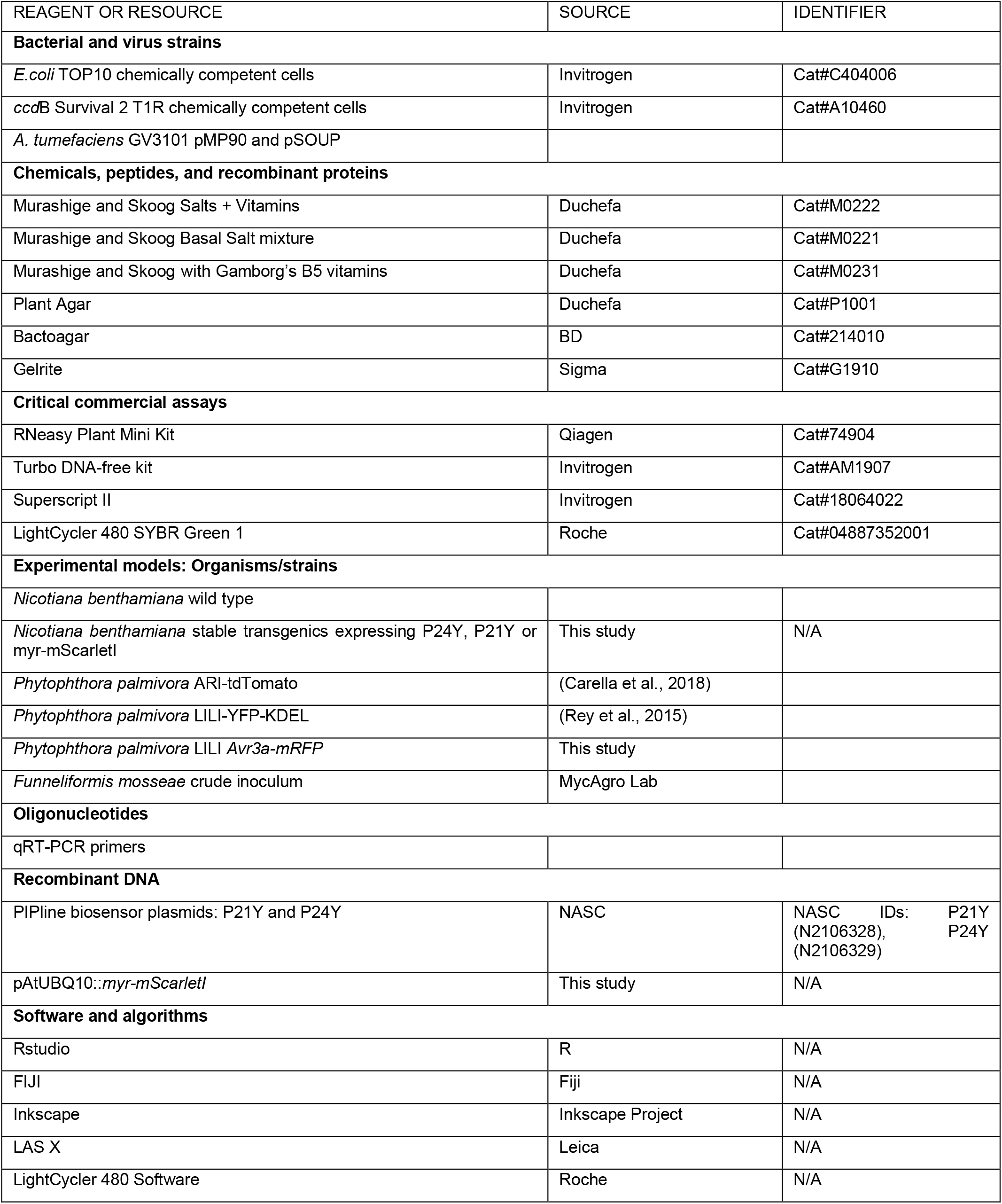

### Plant growth

The *Nicotiana benthamiana* used in this project is a laboratory cultivar obtained from The Sainsbury Lab, Norwich, UK originating from Australia (Bally et al., 2018). Unless otherwise stated, we grew *N. benthamiana* in a glasshouse at 25°C for 16-hr photoperiod.

When grown on plates, we sterilised *N. benthamiana* seeds by incubating for 2mins in 70% EtOH, 10mins in sodium hypochlorite solution (1.8%-4.2% active chlorine), before washing 5x in sterile water. We then grew seeds on ½ MS Plant media consisting of: 0.22% Murashige and Skoog Salts + Vitamins (Duchefa M0222), 0.8% Plant Agar, set to pH 5.7 with KOH. If necessary, we supplemented the media with the relevant selective antibiotics (Kanamycin at 100mg/L or BASTA at 2mg/L). We then grew these plates vertically in a growth chamber set at 24°C with a 16hr photoperiod.

### Phytophthora palmivora growth

*P. palmivora* isolates LILI (reference P16830), initially collected from oil palm in Occidental, Colombia (Torres et al., 2010), and ARI (accession no. P3914, derived from Arizona) are maintained at the Sainsbury Laboratory Cambridge University. We maintained *Phytophthora* strains on V8 vegetable juice agar plates (1.5% agar, 100mg/L G418, 50mg/L Carbenicillin), in a dark Conviron growth cabinet set to 25°C. We collected *P. palmivora* zoospores for infection assays from 7-to 10-day-old plates of *P. palmivora*, as previously described (Le Fevre et al., 2016). Transgenic LILI strains expressing KDEL-YFP (Rey et al., 2015) and ARI expressing tdTomato (Carella et al., 2018) have been previously described. The LILI strain carrying the *Avr3a-mRFP* transgene was obtained via zoospore electroporation (Huitema et al., 2011) of LILI with a pHam34::*PiAvr3a-mRFP* plasmid obtained from (Whisson et al., 2007).

## METHOD DETAILS

### Plasmid construction

We obtained PIP biosensor (Simon et al., 2014) constructs P21Y and P24Y from NASC (NASC stock set # N21056604) as plasmid DNA which we transformed into *A. tumefaciens* strain GV3101-pMP90-pSOUP for use in transient assays and to generate stable transgenics.

We generated pAtUBQ10::*myr-mScarletI* using an entry clone from laboratory stocks (Bonfanti et al., 2023). We recombined this entry clone using an LR reaction into the binary vector pK7m34GW (Karimi et al., 2005) under the control of the AtUBQ10_2000_ promoter with *mNeonGreen* at the C-terminus. Due to the STOP codon in *mScarletI, mNeonGreen* is not translated, but was included as part of the cloning strategy. We maintained empty gateway vectors in OneShot ccdB survival *E*.*coli* cells (Invitrogen). We maintained other plasmids in TOP10 *E*.*coli* cells (Invitrogen). We transformed expression clones into electrocompetent *Agrobacterium tumefaciens* GV3101 containing the helper plasmids pMP90 and pSOUP (Hellens et al., 2000; Koncz et al., 1994).

### Generation of stable transgenics

*N. benthamiana* stable transformation was performed according to Sparkes et al., 2006 with the following modifications. We used leaves from 4wpi wild-type plants infiltrated with *A. tumefaciens* containing the relevant construct. We left leaves on the plant for three days before removal. We incubated leaf discs on shoot-inducing medium (SIM) composed of 1X Murashige and Skoog with Gamborg’s B5 vitamins supplemented with 1% sucrose, 1.5% Bactoagar, 2mg/L 6-benzylaminopurine (BAP), 50mg/L 1-napthaleneacetic acid (NAA), 320mg/L Timentin, 500mg/L of Cefotaxime as well as the construct specific antibiotic (e.g Kanamycin at 100mg/L or BASTA at 2mg/L) at a pH of 5.7. We cut emerging shoots and transferred them to root-inducing medium (RIM), composed of ½ strength Murashige and Skoog Basal Salt mixture supplemented with 0.5% sucrose, 0.25% Gelrite, 50mg/L 1-napthaleneacetic acid (NAA), 320mg/L Timentin, 500mg/L of Cefotaxime as well as the construct specific antibiotic (e.g Kanamycin at 100mg/L or BASTA at 2mg/) at a pH of 5.8. After roots emerged, we transferred plantlets to soil and grew them to seed in a glasshouse at 25°C under 16-hr photoperiod.

### Transient expression

For transient expression of constructs in *N. benthamiana* leaves, we grew *A. tumefaciens* for two days at 28°C in LB media with the appropriate antibiotics. We then resuspended this culture in agroinfiltration medium composed of 10mM of MgCL_2_, 10mM 2-(N-morpholino)ethanesulfonic acid (MES), 200μM acetosyringone at a pH of 5.6. We adjusted the optical density at 600nm (OD_600_) to 0.4 and incubated at room temperature for one hour. We performed agroinfiltrations using a 1ml syringe without a needle on the abaxial side of 4-week-old *N. benthamiana* leaves.

### Confocal imaging

We performed confocal fluorescence microscopy with a Leica SP8 laser-scanning confocal microscope equipped with HyD detectors, a 25x/0.95 numerical aperture (NA) water objective and a 63x/1.2 NA water objective (Leica, Wetzlar, Germany). We used time-gating, with the window set to open at 0.5ns and close at 6ns. When two fluorescent proteins were imaged simultaneously, we used sequential imaging by line to minimise bleed-through. We captured images at a resolution of 1024×1024, with a scan speed of 200Hz and a line average of 4. The imaging settings for various fluorophores and fluorescent stains used in this project are listed in Table S1.

### Root infection

We grew *N. benthamiana* plants on plates for one week before transfer to 12-well plates. We filled each well with 4ml of ½ MS Plant liquid media (0.22% Murashige and Skoog Salts + Vitamins (Duchefa M0222), set to pH 5.7 with KOH). We inoculated with 10,000 spores of *P. palmivora* ARI-td per well. We incubated plates at 25°C under continuous light. We harvested roots for imaging at 2dpi.

### Leaf infection

We detached leaves from 4-week-old *N. benthamiana* stable transgenic plants and placed them abaxial side up on wet paper towels. We inoculated each leaf by application of four 10µl droplets of zoospore suspension (10^5^ zoospores per millilitre). We incubated leaves under constant light at 25°C and imaged at 2 dpi.

For infection following agroinfiltration, we detached leaves after 1 hour and inoculated by applying 10 μl droplets of sterile water or zoospore suspension (10^5^ zoospores per millilitre) at the edge of the infiltrated area. We incubated these leaves at 25°C on wet paper towels for two days before confocal imaging.

### AM colonisation assays

We germinated *N. benthamiana* seeds on Levington F2 compost (ICL, Ipswich, UK) for one week in a growth chamber set at 21°C with a 16hr photoperiod or grew on plates. We then transferred one week old seedlings to 6×4 cellular trays containing sand supplemented with a 1:10 volume of *Funneliformis mosseae* crude inoculum (MycAgro Labs, Bretenière, France). We watered these plants with a low-phosphate Long Ashton nutrient solution (Table S2) three times a week until harvesting (Hewitt, 1966).

We used AMSlides for timelapse imaging of live, AM colonised, *N. benthamiana* plants (McGaley et al., 2024). We grew seeds on plates for one week, then for one week in standard AM colonisation assays (as above) before transferring to AMSlides containing 1:10 volume of *F. mosseae* crude inoculum mixed with sand. We grew the AMSlides in trays lined with damp paper towels and covered with propagator lids in a growth chamber set at 21°C with a 16hr photoperiod. We watered these plants as needed with a low-phosphate Long Ashton nutrient solution (Table S2).

### Co-colonisation assays

We grew *N. benthamiana* plants in the presence of live or autoclaved *F. mosseae* crude inoculum as described above for three weeks. We then harvested these plants, gently washed their roots, and transferred the entire plant to 12-well plates and infected with *P. palmivora* as described above. However, we inoculated these plants with 40,000 spores of *P. palmivora* and harvested roots for imaging or RT-qPCR at 2dpi.

### RNA extraction, cDNA synthesis and RT-qPCR

We collected *N. benthamiana* tissue, flash froze it in liquid nitrogen and ground it using a TissueLyser II (Qiagen). We carried out RNA extraction using RNeasy kits (Qiagen), using buffer RLC and β -mercaptoethanol. To increase the RNA concentration, we reduced the elution volume to 30µl and used the flow through to elute once again. We treated the collected RNA sample with Turbo DNase (Invitrogen) to digest DNA contamination.

We synthesised cDNA using Superscript II (Invitrogen) using 2µg of RNA per sample. We carried out qPCRs on a 384-well LightCycler 480 II machine (Roche), with samples loaded in technical triplicates. Each well contained 2.5µl of 10x diluted cDNA and 7.5µl of PCR mix consisting of 5µl of LightCycler 480 SYBR Green I Master Mix (Roche), 1µl of forward primer (10µM stock), 1µl of reverse primer (10µM stock), and 0.5µl of nuclease-free water. RT-qPCR primers used in this study are listed in Table S3. We used the following cycling protocol: pre-incubation at 95°C for 5mins, followed by 45 cycles of 20s at 95°C, 20s at 60°C, 20s at 72°C and then melting curve analysis. We calculated the Ct values using the second derivative maximum analysis method in the LightCycler 480 software. We calculated ratios using the geometric means of the Ct values for the gene of interest and the reference genes *NbL23* (Liu et al., 2012) or *PpEF1α* (Le Fevre et al., 2016).

### QUANTIFICATION AND STATISTICAL ANALYSIS

We performed statistical analyses of RT-qPCR data in R, carrying out Kruskal-Wallis tests. We reported N numbers and p value cut-offs in the figure legends.

