## Supplementary material for "Mutualist-pathogen co-colonisation modulates phosphoinositide signatures at host intracellular interfaces": Document S1

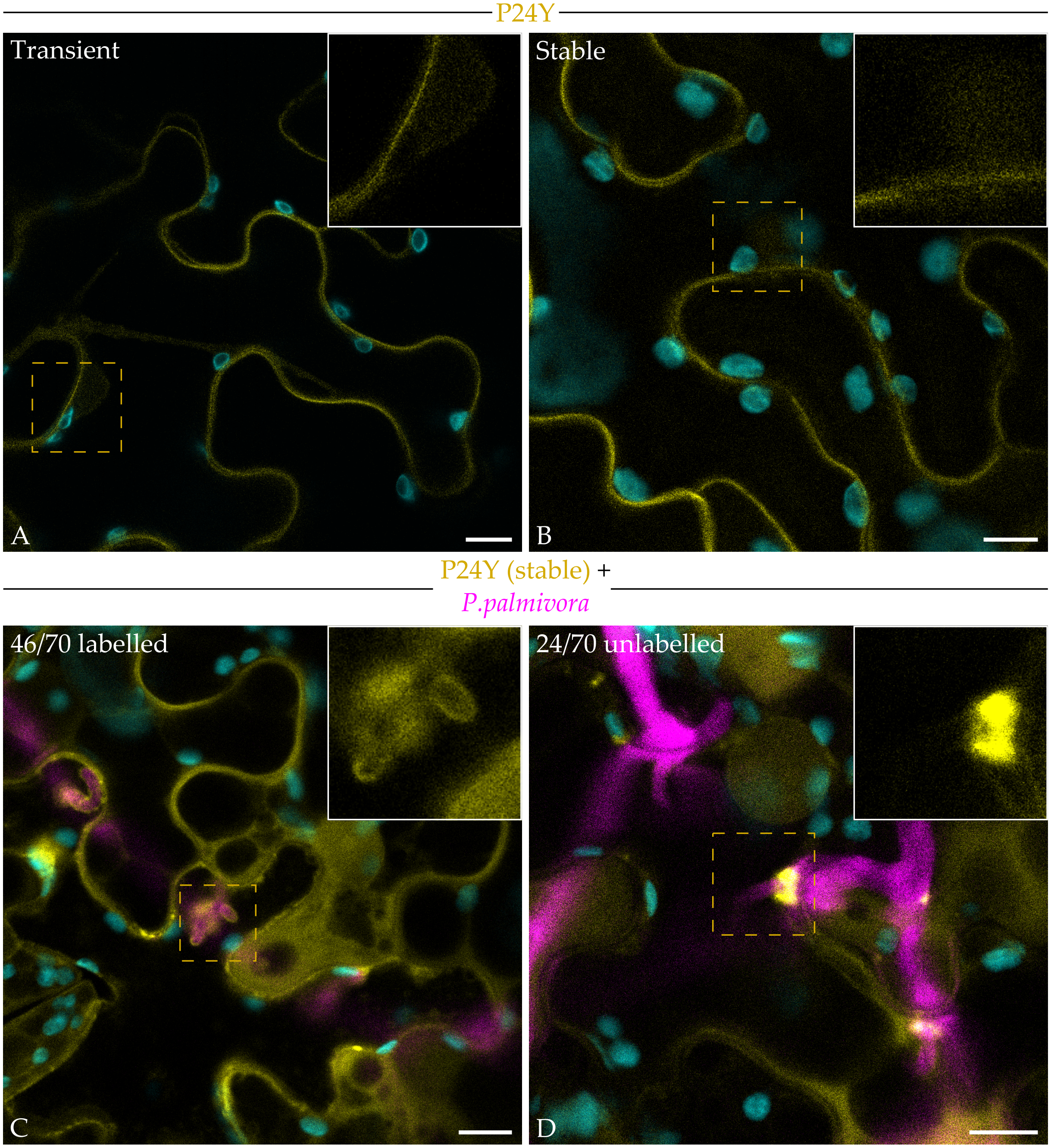


**Figure S1) High-avidity PI(4,5)$P_2$ biosensor P24Y labels *P. palmivora* haustoria in leaves, related to Figure 1.** Confocal imaging of *N. benthamiana* leaf abaxial epidermal cells expressing the PI(4,5)$P_2$ biosensor P24Y (pAtUBQ10::*Citrine-2xPH*$^{PLC}$). **A-B:** P24Y signal in transiently transformed cells (**A**) or stable transgenic cells (**B**). Images are composites of Citrine (yellow) and plastid autofluorescence (cyan) channels. **C-D:** Leaves of stable transgenics were infected with *P. palmivora* strain ARI-tdTomato. Images are composites of Citrine (yellow), tdTomato (magenta) and plastid autofluorescence (cyan) channels. All images are single z-slices. Scale bars = 10µm.

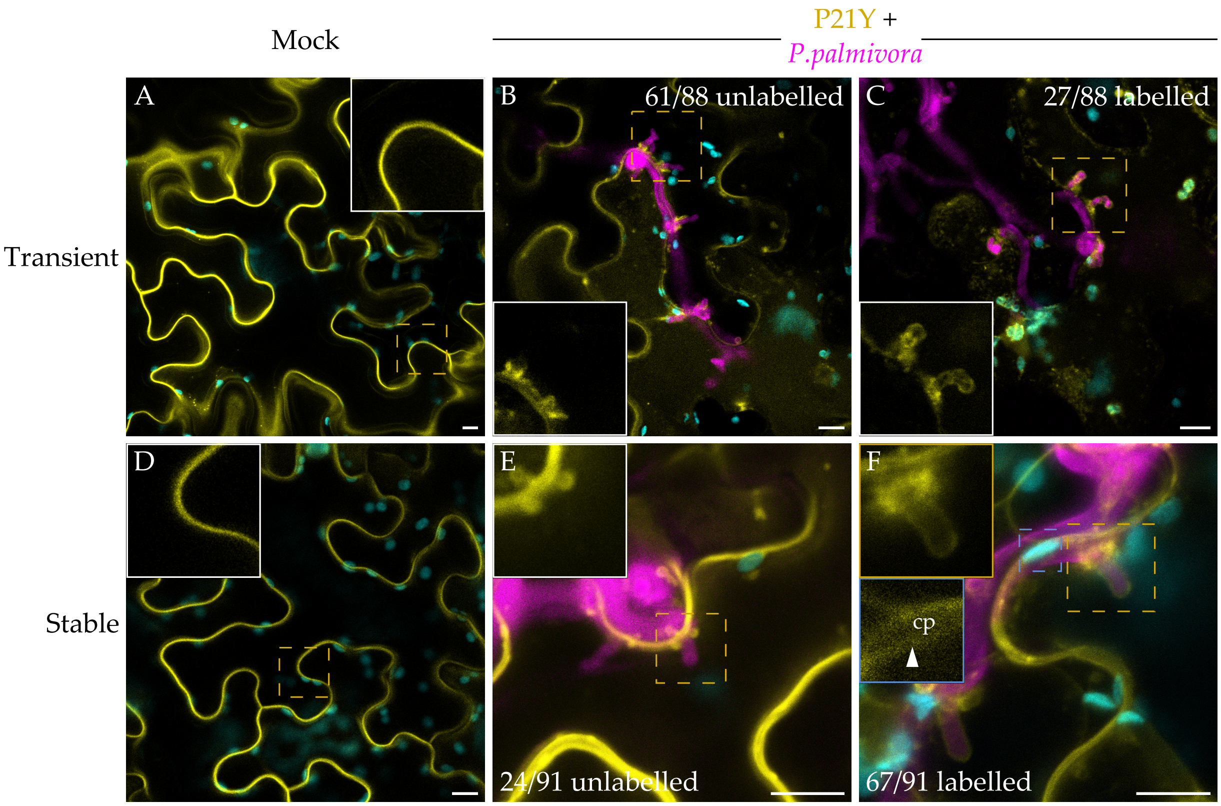


**Figure S2) PI4P biosensor P21Y is sometimes excluded from haustoria in leaves, related to Figure 2**. Confocal imaging of *N. benthamiana* leaf abaxial epidermal cells transiently (**A-C**) or stably (**D-F**) expressing the PI4P biosensor P21Y (pAtUBQ10::*Citrine-2xPH$^{FAPP1}$*). **A, D:** Mock leaves, images are composites of Citrine (yellow) and plastid autofluorescence (cyan) channels. **B-C, E-F:** Leaves were infected with *P. palmivora* strain ARI-tdTomato. Images are composites of Citrine (yellow), tdTomato (magenta) and plastid autofluorescence (cyan) channels. Images A-D are single z-slices, images E-F are maximum intensity z-projections. cp, chloroplast. Scale bars = 10µm.

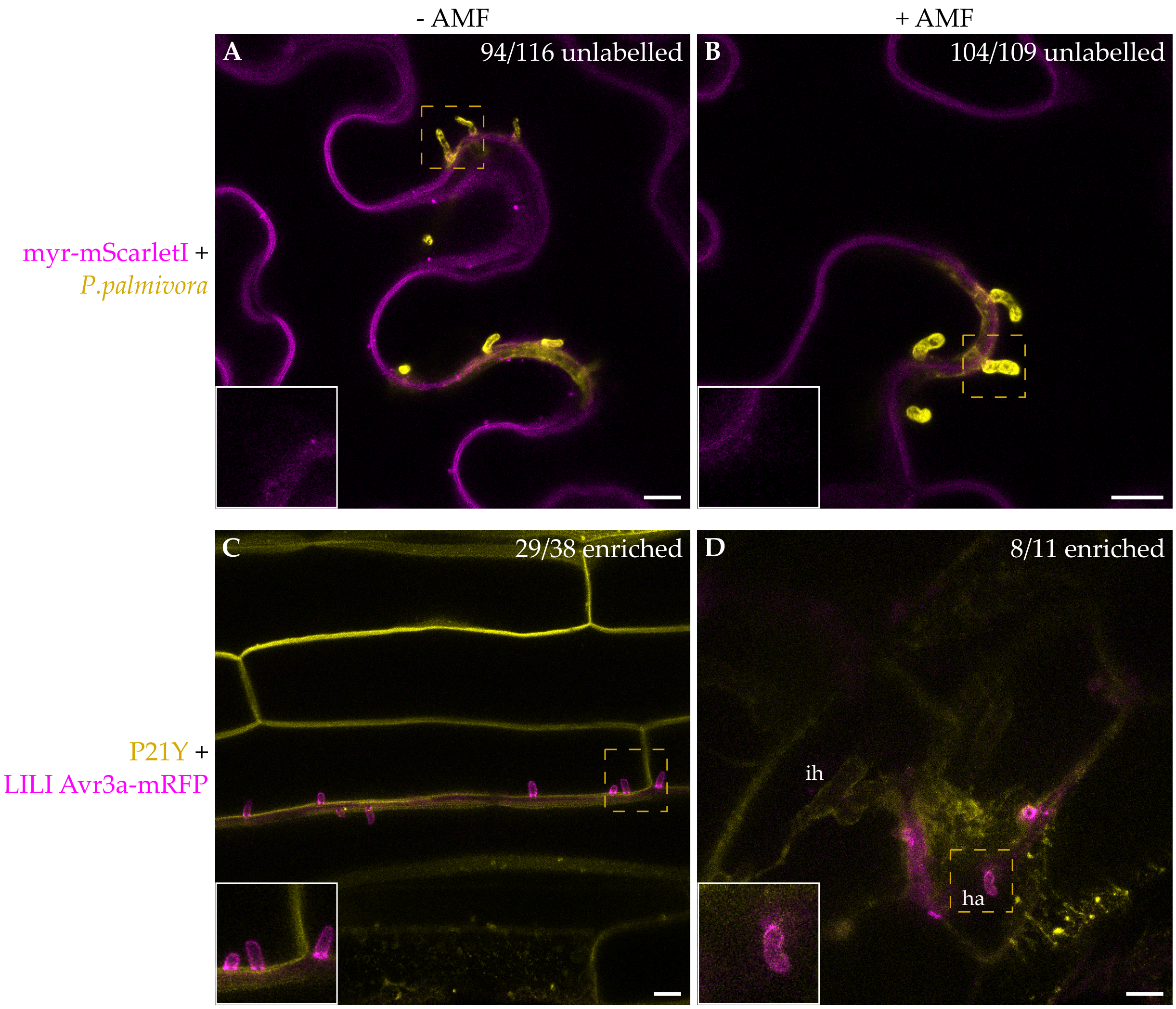


**Figure S3) Myr-mScarletI is excluded from haustoria in leaves of non-mycorrhizal and mycorrhizal plants. Avr3a-mRFP is secreted at haustoria in non-mycorrhizal and mycorrhizal roots. Related to Figure 3. A-B:** *N. benthamiana* stable transgenic leaves expressing pAtUBQ10::*myr-mScarletI* infected with *P. palmivora* strain LILI-YKDEL in the absence (**A**) or presence (**B**) of AM colonisation of the roots. Images are composites of the YFP channel (yellow) and the mScarletI channel (magenta). **C-D:** *N. benthamiana* stable transgenic roots expressing the PI4P biosensor P21Y (pAtUBQ10::*Citrine-2xPH*$^{FAPP1}$) infected with *P. palmivora* strain LILI Avr3a-mRFP in the absence (**C**) or presence (**D**) of AM colonisation. Images are composites of the Citrine channel (yellow) and the mRFP channel (magenta). All images are maximum intensity z-projections. ih, intracellular hyphae; ha, haustoria. Scale bars = 10µm.

**Table S1) Confocal imaging settings, related to Methods.**

| Fluorophore | Excitation Wavelength (nm) | Detection Window (nm) |
|---|---|---|
| Chloroplasts | N/A | 679-713 |
| Citrine | 515 | 520-551 |
| mRFP1 | 584 | 590-636 |
| mScarletI | 569 | 575-625 |
| tdTomato | 555 | 565-600 |
| YFP | 515 | 521-550 |

**Table S2) Low-phosphate Long Ashton nutrient solution, related to Methods.**

| Element | Formula | g/mol | Final mM | Final g/L | 100X Stock g/L |
|---|---|---|---|---|---|
| **Macroelements** | | | | | |
| Potassium nitrate | $KNO_3$ | 101.1032 | 2 | 0.202206 | 20.22 |
| Calcium nitrate tetrahydrate | $Ca(NO_3)_2.4H_2O$ | 236.15 | 2 | 0.472300 | 47.23 |
| Magnesium sulfate heptahydrate | $MgSO_4.7H_2O$ | 246.47 | 1 | 0.246470 | 24.65 |
| Sodium dihydrogen phosphate monohydrate | $NaH_2PO_4.H_2O$ | 137.99 | 0.134 | 0.0184906 | 1.85 |
| **Microelements** | | | | | **2000X Stock g/L** |
| **Solution A** | | | | | |
| Sodium chloride | NaCl | 58.44 | 0.05 | 0.002922 | 5.84 |
| Manganese sulfate monohydrate | $MnSO_4.H_2O$ | 169.02 | 0.005 | 0.000845 | 1.69 |
| Copper(II) sulfate pentahydrate | $CuSO_4.5H_2O$ | 249.69 | 0.0005 | 0.000125 | 0.25 |
| Zinc sulfate heptahydrate | $ZnSO_4.7H_2O$ | 287.56 | 0.001 | 0.000288 | 0.58 |
| Boric acid | $H_3BO_3$ | 61.83 | 0.0165 | 0.001020 | 2.04 |
| Cobalt(II) sulfate heptahydrate | $CoSO_4.7H_2O$ | 281.1 | $5x10^{-5}$ | 0.000014 | 0.03 |
| Nickel(II) sulfate | $NiSO_4$ | 154.76 | $5x10^{-5}$ | 0.000008 | 0.02 |
| **Solution B** | | | | | |
| Sodium Molybdate | $Na_2MoO_4$ | 205.92 | 0.0001 | 0.000021 | 0.04 |
| | | | | | **100X Stock g/L** |
| EDTA iron(III) sodium salt | Na-Fe-EDTA | 367.05 | 0.05 | 0.018353 | 1.84 |

**Table S3) RT-qPCR primers used in this study, related to Methods**

| Target gene | Primer name | Primer sequence | Source |
|---|---|---|---|
| *NbL23* | NbL23-qF | AAGGATGCCGTGAAGAAGATGT | (Liu et al., 2012) |
| | NbL23-qR | GCATCGTAGTCAGGAGTCAACC | |
| *NbPT5b* (Niben101Scf02726g00004.1) | NbPT5b_qF | TTACTACGATCCTGCTACACACG | (Timoneda et al., 2021) |
| | NbPT5b_qR | CAGTTAAAGTACCAACCAAAGCC | |
| *PpEF1α* | PpEF1a-qF | CAAGATCCCGTTCGTGCCTA | (Le Fevre et al., 2016) |
| | PpEF1a-qR | GCGTTCAGGTTGTCAAGAGC | |
